# Continuously perfusable, customisable and matrix-free vasculature on a chip platform

**DOI:** 10.1101/2022.08.02.499348

**Authors:** Francois Chesnais, Jordan Joel, Jonas Hue, Sima Shakib, Lucy Di-Silvio, Agamemnon E. Grigoriadis, Trevor Coward, Lorenzo Veschini

**Affiliations:** Academic centre of reconstructive science, Centre for oral, clinical and translational sciences, Faculty of Dentistry Oral & Craniofacial Sciences, King’s College London, Guy’s Hospital, Great Maze Pond, London SE1 9RT, UK; Centre for craniofacial and regenerative medicine, Faculty of Dentistry Oral & Craniofacial Sciences, King’s College London, Guy’s Hospital, Great Maze Pond, London SE1 9RT, UK; Centre for oral, clinical and translational sciences, Faculty of Dentistry Oral & Craniofacial Sciences, King’s College London, Guy’s Hospital, Great Maze Pond, London SE1 9RT, UK; Centre for gene therapy and regenerative medicine, Faculty of Life Sciences and Medicine, King’s College London, Guy’s Hospital, Great Maze Pond, London SE1 9RT, UK

## Abstract

Creating vascularised cellular environments *in vitro* is a current challenge in tissue engineering and a bottleneck towards developing functional stem cell-derived microtissues for regenerative medicine and basic investigations. Here we have developed a new workflow to manufacture Vasculature on Chip (VoC) systems efficiently, quickly, and inexpensively. We have employed 3D printing for fast-prototyping of bespoke VoC and coupled them with a refined organotypic culture system (OVAA) to grow patent capillaries *in vitro* using tissue-specific endothelial and stromal cells. Furthermore, we have designed and implemented a pocket-size flow driver to establish physiologic perfusive flow throughout our VoC-OVAA with minimal medium use and waste. Using our platform, we have created vascularised microtissues and perfused them at physiologic flow rates for extended times and observed flow-dependent vascular remodelling. Overall, we present for the first time a scalable and customisable system to grow vascularised and perfusable microtissues, a key initial step to grow mature and functional tissues *in vitro*. We envision that this technology will empower fast prototyping and validation of increasingly biomimetic *in vitro* systems, including interconnected multi-tissue systems.

## Introduction

Creating functional human tissues *in vitro* is a current major challenge in both regenerative medicine and basic cell and developmental biology. In the regenerative field, the hope is to create stem cell-derived organ’s functional units as an unlimited source of tissues for transplantation. In the cell and developmental fields, bottom-up creation of tissues is an appealing way to investigate the molecular details of cell differentiation and organogenesis and to reduce/replace animal experimentation. Creating functional tissues *in vitro* requires recapitulating the tissue microenvironment including specific vascular supply which is a currently unmet challenge. Addressing such challenge requires resolving a host of cell biology and engineering problems.

Blood vessels form a spread-out network *in vivo*, supplying organs with nutrients and oxygen, removing waste and enabling a permanent immune surveillance (Potente et al., 2011). Most cells in animal tissues reside within 100 µm from a blood capillary, and angiogenic/angiocrine signalling is key to proper development and homeostasis of an organ’s functional units (Rafii et al., 2016). The cardiovascular system (CVS) is formed by the heart and a hierarchical network of blood vessels from large arteries and veins to tiny capillaries. Engineering such complex system *in vitro* requires taking into account both the differences in cell composition and the biophysical microenvironment experienced by different vessels in the body (Song et al., 2018).

The functional specialisation of blood vessels emerges early during development and vascular cells like endothelial cells (EC), pericytes (PC) and smooth muscle cells (SMC) have heterogeneous transcriptional profiles among different tissues and within the same tissue (Kalucka et al., 2020). It follows that engineering tissue-specific blood vessels necessitates employing such specialised cellular components.

Among all the key factors towards blood vessel maturation and function, flow-generated mechanical forces are essential (Fang et al., 2017; Obi et al., 2009). Applying appropriate mechanical forces *in vitro* requires precisely controlling and regulating the flow of medium or blood substitute entering engineered vessels.

Micro physiological systems have been recently developed to generate and perfuse primitive vascular networks resembling that observable during initial phases of wound healing (Chen et al., 2017; Hasan et al., 2014; Shin et al., 2012; van Dijk et al., 2020). These systems leverage spontaneous or guided endothelial cells (EC) self-assembly within 3D matrices like fibrin or collagen. One advantage of these systems is that perfusable vascular-like tubes or even networks can be easily created in 2-3 days, however these networks are not reminiscent of mature, fully functional capillaries because hydrogels cannot support vessel maturation and remodelling in absence of matrix producing cells. Furthermore, continuous perfusion (> 24h) has not been extensively demonstrated due to limitations in creating secure connections between the vasculature and the flow generator (e.g., peristaltic or syringe pumps). In fact, perfusion has been mostly demonstrated using gravity-driven flow which is scarcely controllable and not maintainable for periods longer than a few hours. Finally, current systems manufactured by SU-8 photolithography (Chen et al., 2017; Hasan et al., 2014; Song et al., 2018; Wang et al., 2016; Zheng et al., 2012) are relatively fixed in design and expensive to manufacture thus, they are not amenable to fast-prototyping of multicomponent systems which are key for future development of multiple connected organs on chip (Ronaldson-Bouchard et al., 2022).

To overcome all these challenges, we have developed a new manufacturing workflow for inexpensive and fast prototyping of custom microfluidic devices by 3D printing followed by PDMS soft lithography. Using this workflow, we have designed, manufactured, and tested a new microfluidic lab-on-chip (LoC) to grow capillary-like structures including secure world to LoC connections for their perfusion. Furthermore, we have refined a matrix free system to robustly grow patent capillary networks by organotypic culture of EC and matrix-producing stromal cells (OVAA) and adapted it to our LoC. Finally, we have created a “plug-and-play” platform for chip perfusion which can fit directly in a cell culture incubator enabling maintenance of optimal medium temperature and minimal waste. We have validated our system under different flow conditions and demonstrated tissue perfusion and capillary remodelling in response to continuous flow.

Overall, we present for the first time a scalable and customisable micro physiological system to grow biomimetic, vascularised and perfusable microtissues as a key initial step to enable growing mature and functional tissues *in vitro*.

## Results

### 3D printing-based manufacture enables rapid prototyping of LoC devices

Creating tissue-specific perfusable vascular networks (Vasculature on Chip, VoC) requires flexibility in design as different tissues have different types of blood vessels in terms of dimensions, flow rate, type of endothelial barrier, mural cells coverage and ECM (Aird, 2007; Augustin and Koh, 2017). Current techniques to manufacture microfluidic chips are laborious and expensive (Chen et al., 2017) therefore, to build our VoC prototype we developed a workflow based on 3D Computer Assisted Design (3D CAD) and 3D printing (3DP) enabling fast prototyping of easily modifiable and low-cost devices (Fig.1).

**Figure 1:**
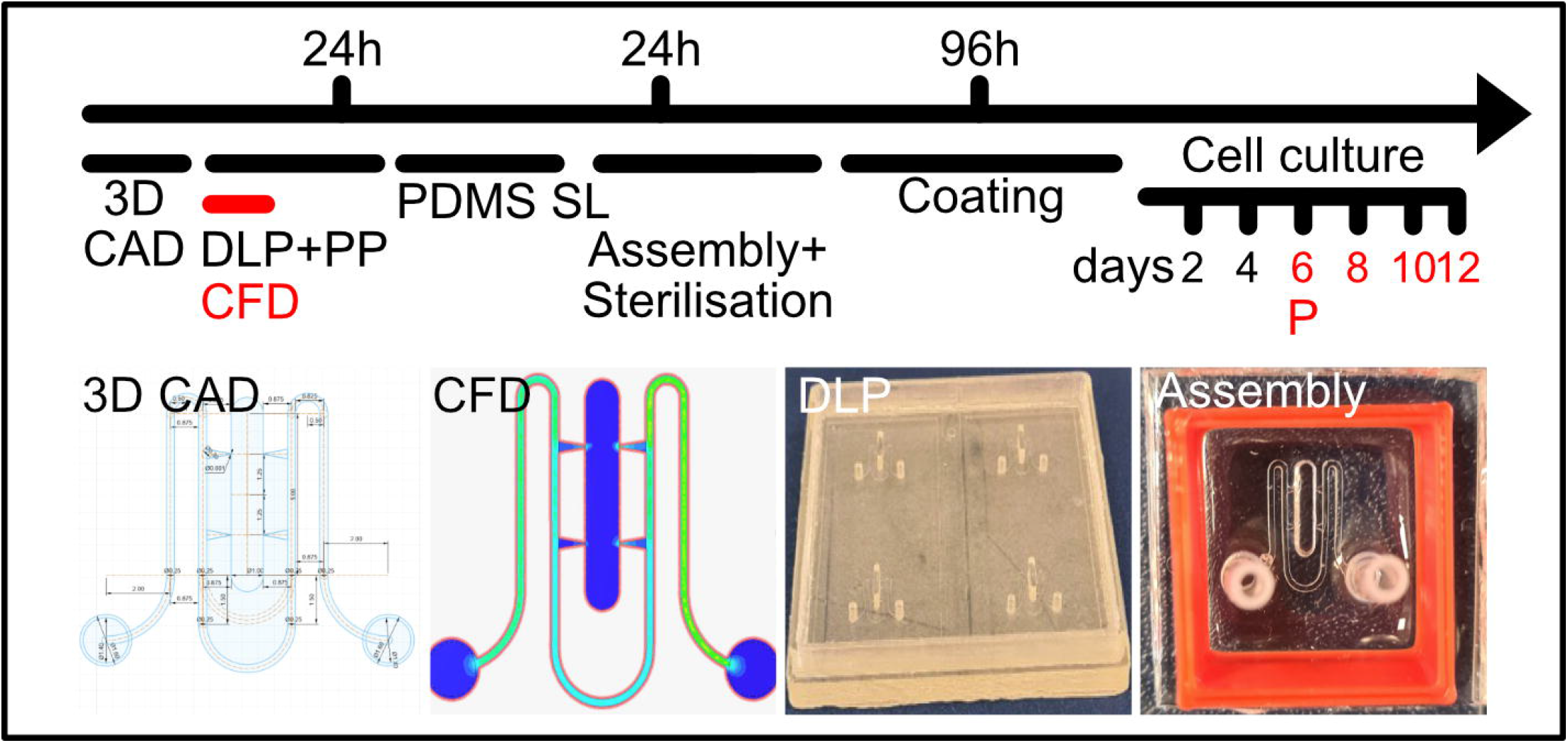
LoC fabrication timeline. Steps include 3D Computer Assisted Design (CAD), mould 3DP by Digital Light Processing (DLP), post-processing (PP) and Computational Flow dynamics (CFD) that can be finished in a day. On day 2 the LoC is manufactured by PDMS soft lithography (SL) and assembled on a glass slide including embedded tubing and a 3D-printed medium reservoir. On day 3 and after sterilisation, the LoC can be coated/functionalised and used in cell culture experiments.

3D CAD/3DP enables creation of complex objects (for example, features with different heights) surpassing in terms of flexibility the capabilities of SU-8 photolithography. Furthermore, 3D CAD designs can be readily analysed via computational flow dynamics (CFD) software. Our 3D CAD/3DP workflow allowed us to create secure VoC-to-world connections without dead volume which is key to achieving stable flow through the VoC (Fig. 2A). We also employed fused deposition modelling (FDM) 3DP to create custom “infrastructures” to facilitate experimental procedures (Figs. 1 and 2B-C). The whole workflow from design to experiment, including printing of all appliances and CFD simulations can be completed in 3 days (Fig. 1).

**Figure 2:**
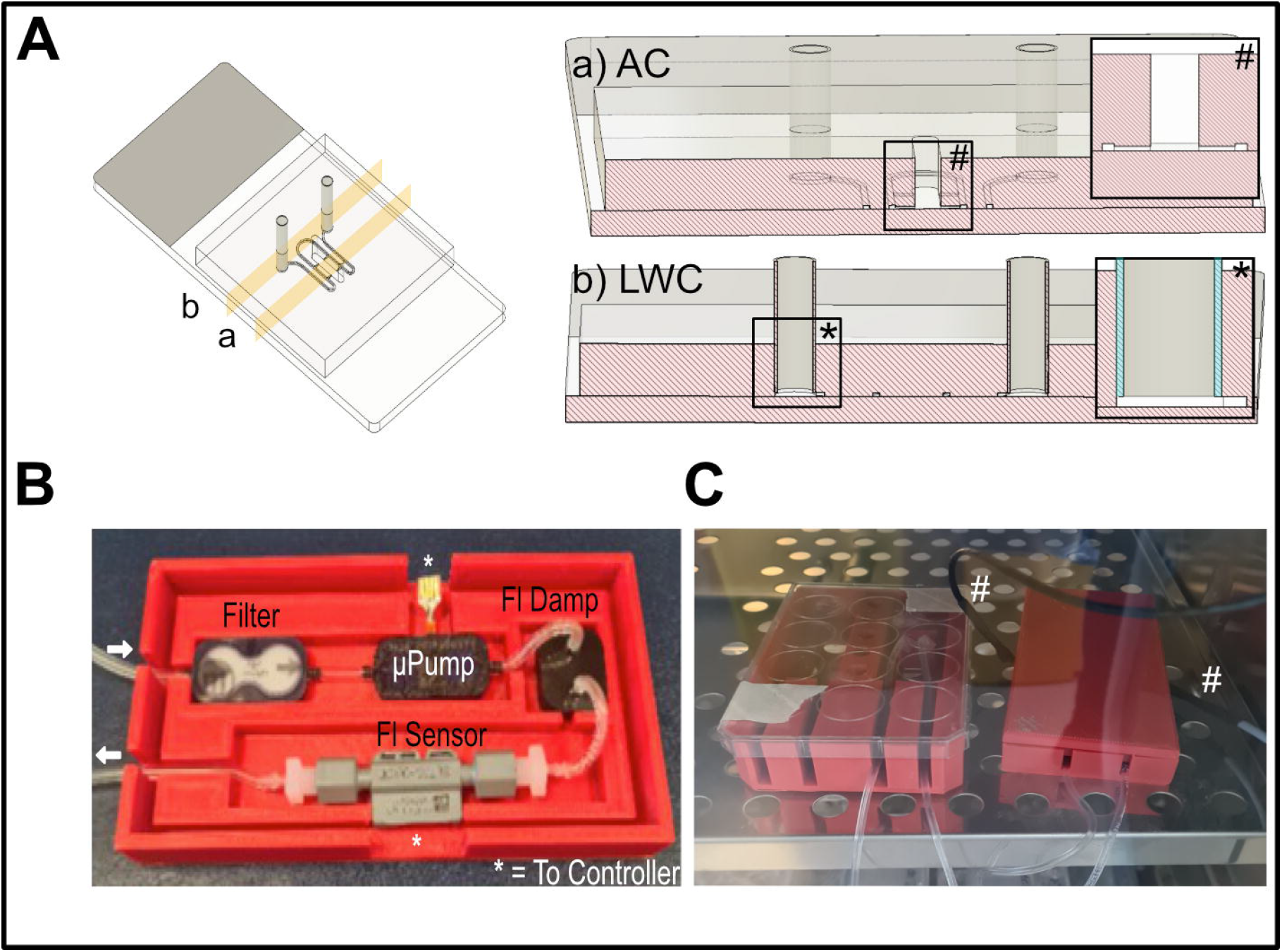
VoC design. A) 3D rendering of the VoC design and corresponding cut-plane analysis at indicated locations (a, b). Cut plane across the Anastomosis Channel (AC, a) highlights the smaller section of the AC in comparison to the main channels. Cut plane across the LOC to World Connection (b, LWC) channel highlights the tight coupling between the chip and the silicone tubing (blue shading in inset) which prevents creation of dead volume and allows predictable flow throughout the chip. B) Pocket-size microfluidic drive in its 3D printed cassette illustrating assembly of the system. C) Microfluidic driver (right) connected to the culture cassette (and chip, left) and positioned in a standard cell culture incubator. The black/grey cables visible on the right (# sign) are the only connection outside the incubator, therefore all medium circulating in the system is exposed to a constant environment.

### Custom design and assembly of VoC platform including a compact perfusion system

To design our VoC, we focussed on two main goals, 1) to mimic physiologic flow rates in pre-capillary and capillary networks and 2) to enable continuous and stable perfusion. To achieve the first goal, we created a two-compartment device. The first is an open-top culture chamber to develop organotypic cultures, and the second compartment is the main channel carrying flow through the chip (Fig. 2Aa). The main channels were designed to mimic afferent arterioles (aA) and efferent venules (eV) of a tissue functional unit (TFU, e.g., a sector of the retina or dermis). Different potential designs were considered (SFig. 1), including separate or connected afferent and efferent channels (SFig. 1A i-v) and different geometries for the connections (SFig. 1B i-iii). We selected the design shown in Fig. 1A where the aA and eV are connected in a loop carrying most of the total flow going through the system. The geometry of these channels is fixed therefore the dynamics of fluid flow in the chip can be predicted using Computational Fluid Dynamics simulations (CFD). The aA/eV system connects to the culture chamber through accesses which were shaped to mimic branching points in arterioles/venules found in vivo (e.g., retina). In line with the literature in the field we aimed to achieve flow rates (FR) of 100-500 µl/min (main channels) leading to a FR of 10-50 µl/min across the connections which are like that of aA and eV *in vivo* in brain capillaries (Chaigneau et al., 2019, 2003).

Continuous perfusion (>24h) of chips containing living cells requires producing appropriate FR using cell culture medium which must be maintained at a constant temperature and in a controlled environment. To achieve these goals, we employed a miniature fluidic driver system which can be assembled and fitted directly in a cell culture incubator (Figs. 2B and C). Fig. 2B shows the flow driver system assembly positioned in its cassette. The flow is driven by a micro-peristaltic pump capable of producing 10-1000 µl/min FR. Flow is stabilised by a flow dampener and FR is measured by a flow sensor. Both the pump and the flow sensor are connected to a controller creating a feedback loop which adjusts pump output to maintain a stable FR.

### CFD analysis predicts biomimetic flow in our VoC design

To optimise the design of our VoC, we examined our candidates (SFig. 1) by CFD and iteratively refined our design aiming to achieve balanced flow between inlet and outlet (in-flow = out-flow) as well as a pressure drop between aA and eV. In this way, the total volume of medium in the system remains fixed while creating flow across the open-top central chamber. Fig. 3 shows CFD analysis of our current design (analysis parameters are listed in methods). Fig. 3A demonstrates that CFD particle analysis predicts establishment of stable flow without turbulences in the system at the indicated FR. Particle speed analysis (FR=100 µl/min) predicted values of ∼10 mm/s within the central chamber and ∼ 100 mm/s in the aV and eV sections. These values are compatible with that observed for red blood cells in brain capillaries and arterioles (4-10 mm/s and 20-30 mm/s respectively) (Chaigneau et al., 2019). Simulations using FR of 1000 µl/min predicted formation of vortices in the LoC outlet, as turbulence could create bubbles in the medium, in experiments we employed maximum FR of 500 µl/min. Fig. 3B shows cut plane analysis (plane at 50 µm distance from the chip base, corresponding to lines in Fig. 3D i-iii) displaying pressure in the different sections of the chip and demonstrating a pressure drop between the aA and eV sides (red and blue respectively) under both FR tested. Fig. 3C shows a zoomed-in representation of the cut-plane analysis (corresponding to the red box in Fig. 3B) including predicted flow speed and directions across a section of the central well. Fig. 3D shows a cut-plane analysis across planes perpendicular to points indicated in the 3D diagram (cross-section of the channels at corresponding points, FR=100µl/min). The colour scale indicates flow speed in different points of the cross-section and the corresponding total volumetric flow rates (VFR).

**Figure 3:**
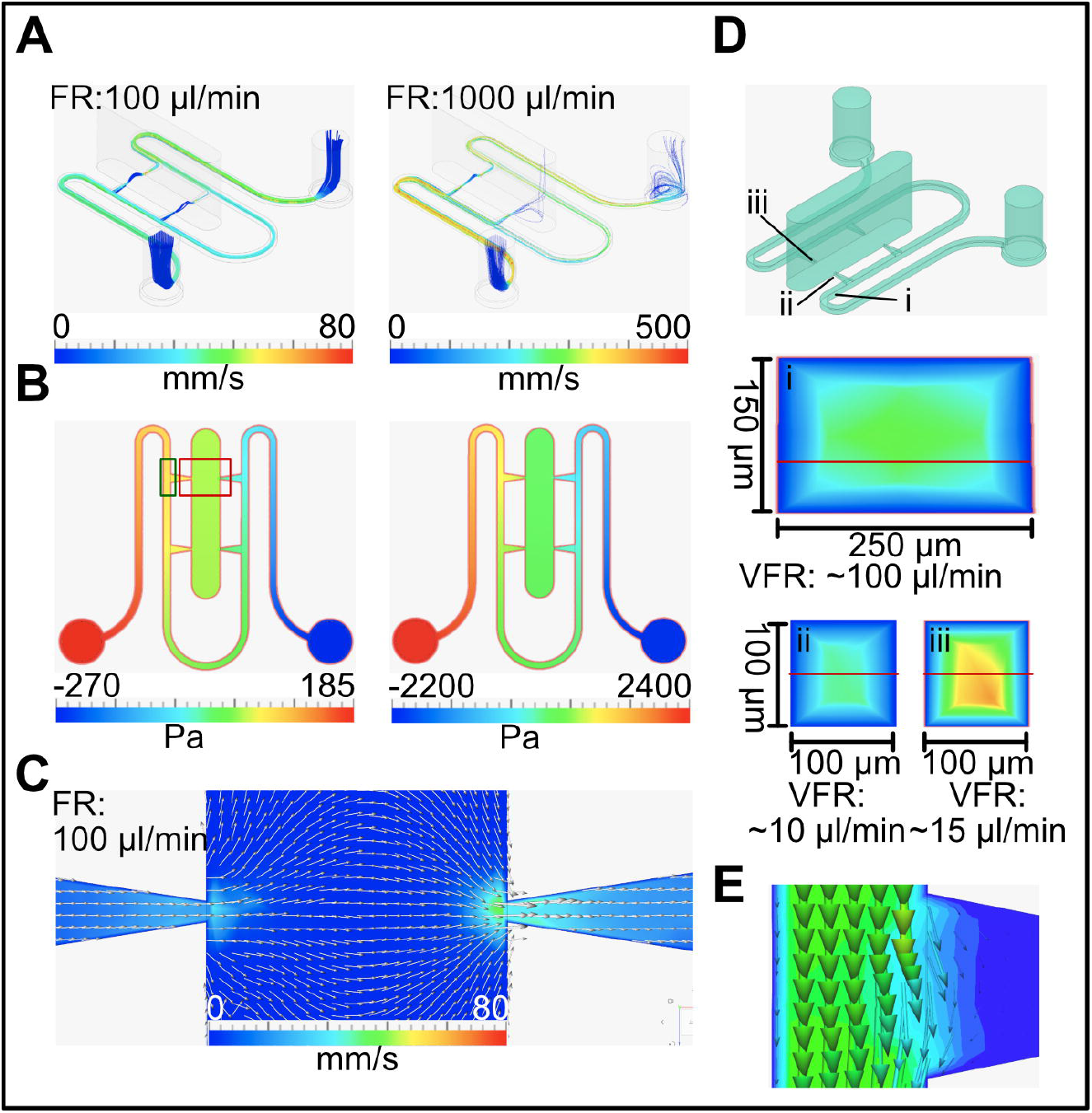
CFD analysis of the VoC design. A) Particle speed analysis at indicated FR. Particles trajectories are indicated by coloured cylinders, colour scale indicates particle speed at each point. B) Cut plane analysis across the X/Y plane indicated in D i-iii and at indicated FR, colour scale indicates pressure at each point in the chip. Pressure in the central chamber is ∼0 (atmospheric pressure, green colour). Red and green coloured boxes indicate locations of analyses illustrated in C and E respectively. C) Vector analysis across the X/Y plane indicated in D i-iii and at indicated FR. Colour scale indicates flow speed at each point, vectors indicate relative speed (intensity) and direction of the flow in a grid of points in the chip section. D) 3D rendering of the VoC geometry indicating regions highlighted in i-iii. D i-iii are cut plane sections at indicated points and with direction X/Z (i) and Y/Z (ii and iii). Colour code indicates flow speed at each point in the cross-sections (colour scale equivalent to A and C). Total Volumetric Flow Rate (VFR) across the sections is as indicated. E) Vector analysis of a simulation where flow through the central chamber was blocked. Analysis corresponds to a section of the chip indicated in B (green box) and across the X/Y plane indicated in D. Colour scale (both vectors and background) correspond to flow speed in the section. Vectors directions correspond to flow direction in a grid of points in the section.

Overall, our CFD analysis demonstrates that the design of our LoC produces desired biomimetic flow rates and directions. In our intended experimental workflow, the flow is constrained differently at different phases of the experiment. Especially, during the first phases, the central chamber is not accessible to flow because the connections are purposedly closed by cells. To assess the behaviour of our design in these conditions, further simulations were performed where the flow access to the central chamber was blocked. Fig. 3E shows CFD analysis of our design under these conditions, demonstrating variation in the flow directions creating an interstitial-like flow at the interfaces between channels and cells. Under these conditions the flow velocity in the central chamber and most of the connection was ∼0.

### Continuous perfusion of self-assembled vascular networks in hydrogels

To validate our system for continuous perfusion we performed preliminary experiments using perfusable vascular networks created in fibrin hydrogels. We encapsulated Endothelial Cells (EC) in fibrin hydrogels where ECs self-assemble to form perfusable structures in approximately 72h (Fig. 4).

**Figure 4:**
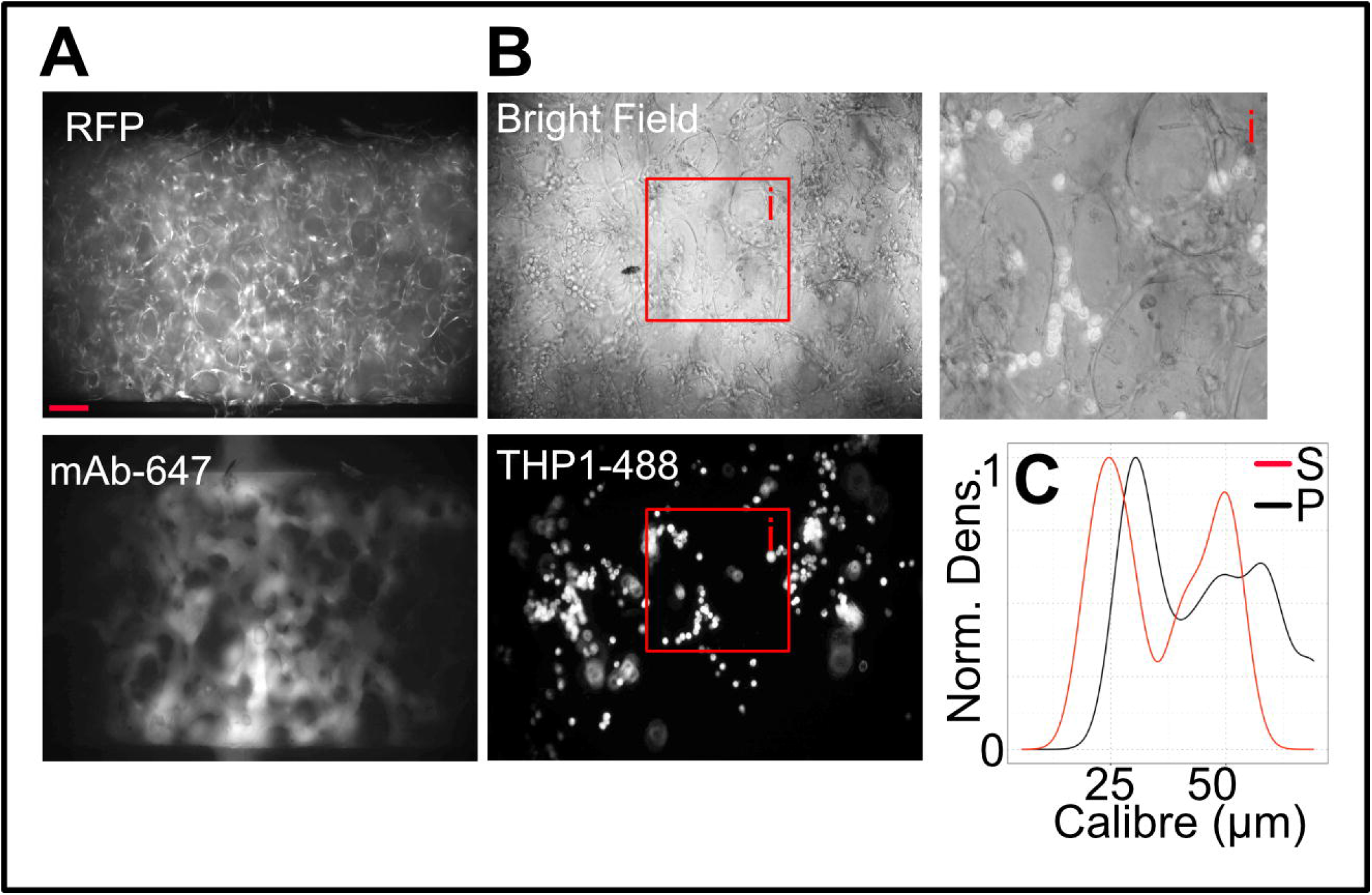
Perfusion of self-assembled network in hydrogels. A) Photograph of the vascular network formed in a fibrin hydrogel by HUVEC-RFP after 72h in culture. Images represent a snapshot of the HUVEC-RFP signal on top and the mAb-647 signal on the bottom during continuous perfusion of the antibody at 150μL/min. Scale bar: 200μm B) Photograph of the vascular network formed in a fibrin hydrogel by HUVEC after 72h of culture. The network was perfused with THP-1 monocytes stained with CellBrite® Cytoplasmic Membrane dye Green (Biotium, UK) and a snapshot was taken for brightfield signal and green, fluorescent signal. A crop (i) shows the close interaction of THP-1 in perfused capillaries. C) Quantification of microvessel width (in μm) formed in fibrin hydrogels before (red trace) or after perfusion (black trace).

To demonstrate perfusion of the vascular structures in our VoC we inoculated a bolus of fluorescently labelled (Alexa 647) secondary anti mouse-specific antibody (mAb-647). Fig. 4A shows fluorescence images relative to a representative experiment 1s after the mAb-647 has flowed through the main channels (FR =150uL/min). The top panel corresponds to an RFP signal of RFP-tagged HUVEC, the lower panel corresponds to signal from the mAb-647 demonstrating that the vascular-like structures are fully perfused. The same microscopic field returned to very low signal after 5s of perfusion demonstrating that the mAb didn’t leak through the network in the timeframe of this experiment (60s). Inspection of the lumen of the structures as highlighted by the mAb-647 perfusion demonstrates that they lack hierarchical organisation and that they are reminiscent of sinusoidal/immature vasculature as that found during early phases of wound healing. To further demonstrate effective perfusion, we performed an experiment where we inoculated a bolus of THP-1 monocytes, reasoning that monocyte should adhere to activated EC such as those in engineered vascular networks. As expected, THP-1 monocytes entered the engineered vascular network and several adhered to the ECs (Fig. 4B main and zoomed in view, i). Finally, to estimate the structure of these networks we measured the calibre of several structures in independent experiments (Fig. 4C). Quantification of structures with or without active perfusion demonstrates that 1) in average vascular-like structures in fibrin hydrogels have calibres of 25-80µm (5-10 times bigger than small capillaries observed *in vivo*) and 2) that perfusion further enlarges these structures demonstrating creation of positive pressure within the network.

### Organotypic culture of human EC and HDF allows creating biomimetic capillary networks

To overcome the limitation of hydrogel based systems, we have refined an existing organotypic co-culture assay (Organotypic Vasculo-/Angio-genesis Assay, OVAA) (Bishop, 1999; Wei et al., 2021) to create biomimetic vascularised microtissues. Previous work has shown the value of the OVAA to create patent capillary networks using HUVEC and human dermal fibroblast (HDF) in a two-week timeframe. Here we have optimised the OVAA to generate capillary networks using different organ specific EC and stroma. Fig. 5A shows representative low magnification images of our optimised OVAA employing different endothelial cells in control conditions or under VEGF stimulation (after CD31 staining). Qualitative and quantitative analyses of microvascular networks generated by the OVAA demonstrate that the system faithfully recapitulates the reported effect of VEGF on angiogenesis and highlights striking differences in the intrinsic angiogenic potential of EC from different vascular beds. Fig. 5B shows high-resolution images of the same capillary-like structures highlighting how the OVAA allows investigating key angiogenic processes such as tip cell selection, vascular branching, intussusceptive angiogenesis, and generation of filopodia.

**Figure 5:**
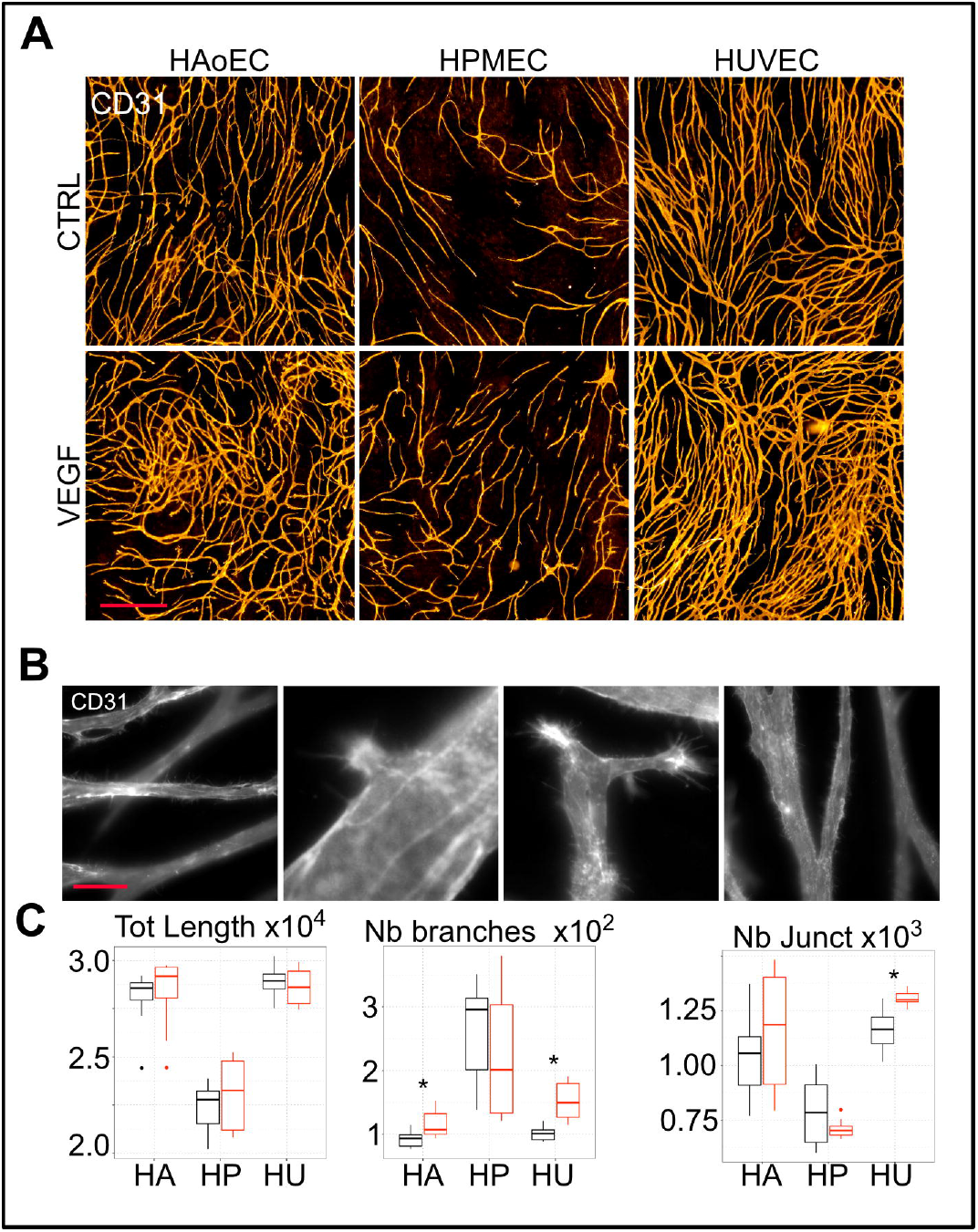
Endothelial/fibroblast co-culture vasculogenesis/angiogenesis assay (OVAA). A) Photographs of vascular networks formed by human aortic endothelial cells (HAoEC), human pulmonary endothelial cells (HPMEC) and human umbilical vein endothelial cells (HUVEC) in co-culture with human dermal fibroblasts (HDF) for 14 days and immunostained for CD31 (PECAM1). Panels show representative images of the different EC lines angiogenic potential in control conditions (EGMV2 medium) or after treatment with 50ng/mL VEGFA. Scale bar: 200μm B) Representative images of vascular processes such as tip cell selection, vascular branching, intussusceptive angiogenesis and filopodia in the vascular networks formed by HUVECs in OVAA, stained for CD31 after 14 days and images at 63X magnification. Scale bar: 50μm C) Quantification of the vascular network complexity in the OVAA with FIJI and the AngioAnalyser plugin. Parameters such as total network length, number of branches and number of junctions have been quantified in control conditions (black) or upon VEGF treatment (red) for the 3 different EC lines in 4 separate experiments and for 2 donors for each EC line.

As we previously reported (Chesnais et al., 2022), ECs isolated from different vascular beds are highly heterogeneous at the population and at the single cell level. This also resulted in appreciable differences in their angiogenic potential as highlighted in Fig. 5C. Although, all 3 lines responded to VEGF treatment as expected (i.e., an increase of the network complexity), they had different intrinsic angiogenic potential under similar experimental conditions. HUVEC and HAoEC had a high proliferative and angiogenic phenotype, forming a highly interconnected network whereas the HPMEC only formed few thin vascular structures.

Overall, using our optimised OVAA, we were able to grow capillaries using combinations of different primary EC (HUVEC, HPMEC, HAoEC) and ECM secreting cells such as HDFs or mesenchymal stem cells (MSC, SFig. 2C); thus, OVAA enables recapitulating different vascularised tissue microenvironments.

### OVAA capillaries in VoC are perfusable and remodel over time in response to flow

As shown above, hydrogel systems are convenient to create and perfuse self-assembled vascular networks, however, they do not allow vascular maturation and remodelling. To overcome this key issue, we combined our OVAA and VoC perfusion system as detailed in methods to generate biomimetic vascular networks which can remodel over time in response to flow.

To this end we used our VoC and created a compartment with “large calibre vessels” formed within the main channels of the chip and a connected compartment (open top central chamber) where OVAA is used to form a capillary network. Fig. 6A demonstrates that after 7-12 days in culture, the ECs formed a densely interconnected capillary network in the central well (see also SFig. 2 A and B).

**Figure 6:**
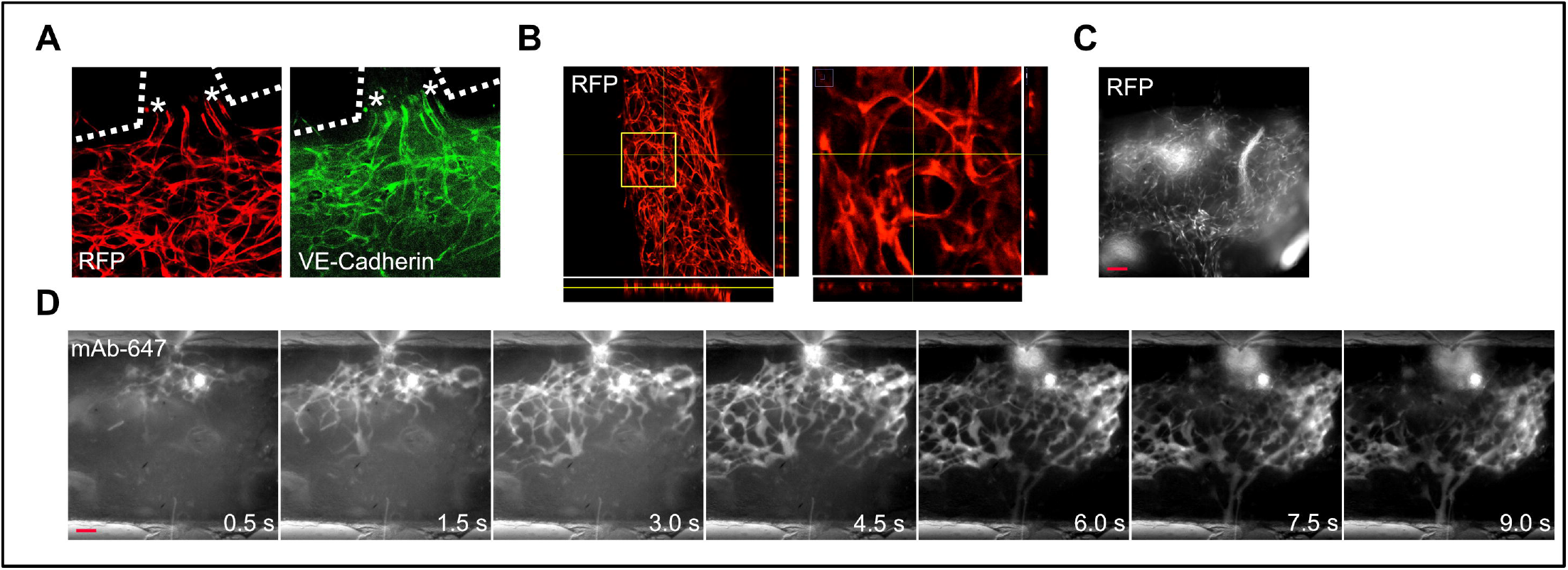
Matrix-free formation of perfusable vascular networks. A) Photographs of HUVEC-RFP signal and VE-cadherin staining in the VoC device after 3 days of perfusion, at the junction between the central well and the side channels. B) Confocal images of HUVEC-RFP vascular neworks formed in the VoC after 7 days of culture. The 3D-projection shows the complexity of the network and the lumen formed by ECs in the micro-tissue. C) Snapshot of the vascular network formed by HUVEC-RFP in culture in the right side of the VoC system and perfused for 10 days. Scale bar: 200μm D) Time series of the same network perfused with mAb-647 (10μg/mL) at a flow rate of 500μL/min with images taken every second. Scale bar: 200μm.

Fig. 6B shows a z projection reconstruction of a 100µm confocal z-stack through a vascular network created with HUVEC and HDF which has been perfused for 1d before fixation and immuno-staining.

Imaging of RFP and VE-Cadherin shows the formation of a densely interconnected EC network including vessels through the intersection channel (* sign in Fig. 6A). Fig. 6B shows orthogonal views projections of the same confocal stack and highlighting capillary structures disposition in multiple interconnected layers. Inspection of high magnification orthogonal Z projections confirmed that the structures possessed a patent lumen (Fig. 6B right panel, also see SFig. 3).

To demonstrate that our microvascular structures are perfused, we inoculated a bolus of mAb-647 (FR = 500 µl/min) and recorded fluorescence images of the central chamber of our VoC.

Fig. 6C shows the network formed by HUVEC-RFP in the OVAA-VoC after 10 days of perfusion. Fig. 6D shows a representative time-series sequence of recordings demonstrating the mAb-647 bolus can transit through the capillaries in ∼10s, differently from parallel experiments using fibrin gels (Fig. 5) where the mAb transited completely in less than 5s (and under lower FR = 150µl/min). Further inspection of the time-lapse recording and the capillary structures highlighted by the A647 fluorescence clearly suggests establishment of preferential flow directions through the networks (the central portion of the network is perfused and emptied faster than the lateral portions) and absence of leakage. Furthermore, OVAA-VoC generated capillary networks with a clear hierarchical structure in comparison to enlarged sinusoidal ones observed in the fibrin system. Overall, these data demonstrate that our OVAA-VoC combination enables the creation of vascularised and continuously perfusable microtissues.

We qualitatively observed that OVAA-VoC networks under long-term perfusion remodelled over time creating organised structures as observed *in vivo*. Fig. 7A shows representative images of the same portion of an OVAA-VoC network after one or ten days of perfusion clearly demonstrating remodelling and maturation of the capillary structures in response to continuous flow. To quantify this effect, we measured the calibre of several randomly chosen capillaries at one, three and nine days upon perfusion in three independent experiments. Results of this analysis (Fig. 7B) showed that the capillaries created with OVAA-VoC are smaller than those generated in fibrin (red trace in Fig. 7A and 4C, ∼5-40 µm vs 25-80 µm respectively) and that their size increases in response to positive pressure generated by flow. Density distributions of vessel calibre counts also highlighted that over time the network remodelled in such way to progressively generate a hierarchy of larger, medium sized and smaller vessels as highlighted by the three distinct peaks in Fig. 7B panels 3d and especially 9d. Since parallel experiments in absence of perfusion did not show similar reorganisation of the capillary structures, we hypothesize that mechanical forces due to perfusive flow in our system are necessary and sufficient to induce initial network remodelling and establishment of a hierarchical architecture.

**Figure 7:**
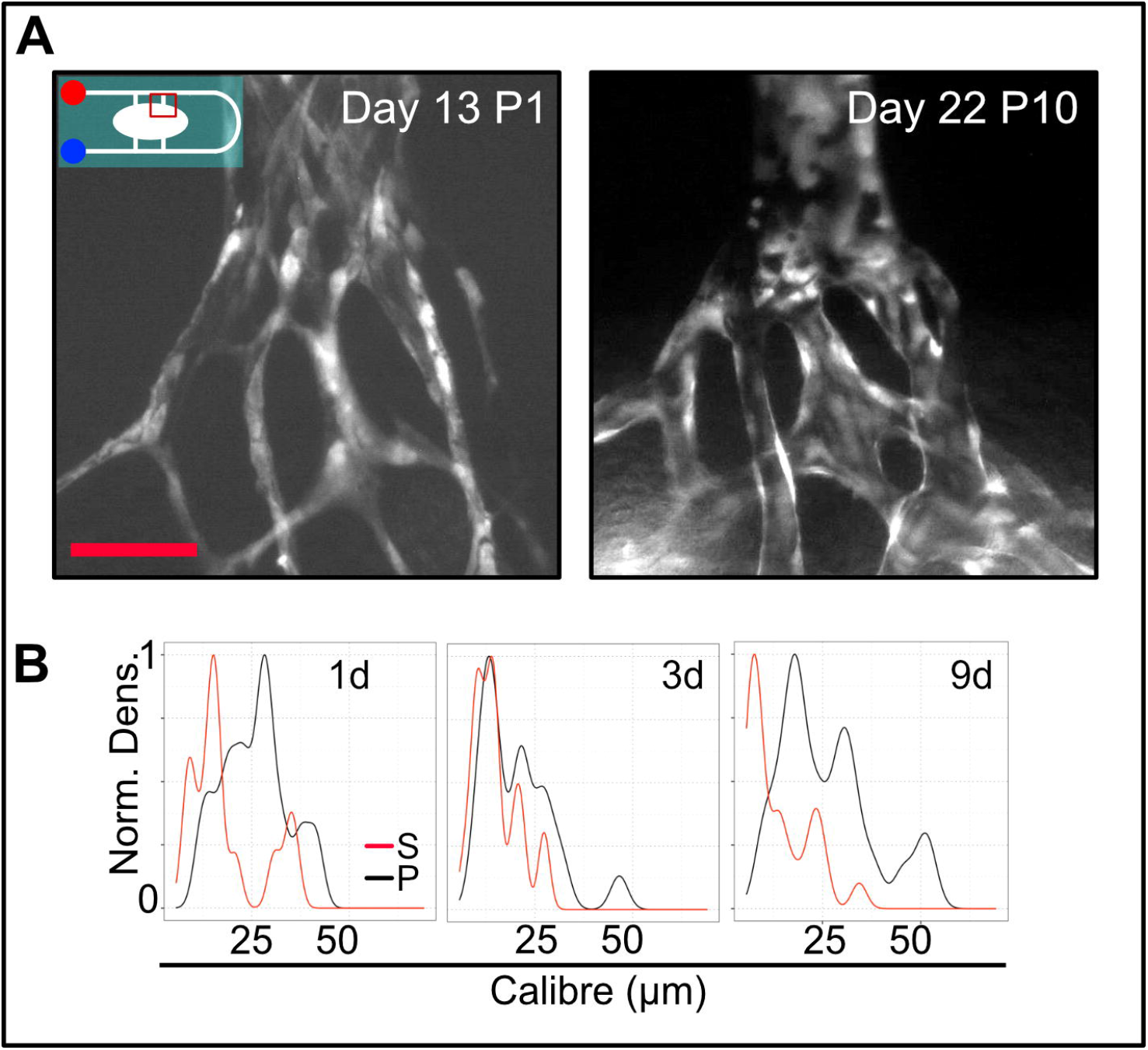
Vascular network remodelling over time in response to flow. A) Photographs of HUVEC-RFP microvessels formed in the VoC corresponding to the region highlighted in red box in schematics, after 13 days of culture (1 day of perfusion) and after 22 days of culture (10 days of perfusion) at 500μL/min. Scale bar: 50μm B) Quantification of the blood vessel structure’s calibre in the the VoC system after 1, 3 or 9 days of perfusion (black line) or without perfusion (red line) for 3 distinct experiments.

Our optimised OVAA-VoC system creates a relatively thick (5-10 cells) tissue in the central well. We observed that when we stopped perfusion in a fully perfused OVAA-VoC the small calibre vessels collapsed (red trace Fig. 7B) suggesting presence of elastic forces within the tissue. This observation is also compatible with the need of high FR (in comparison to fibrin) to achieve perfusion and with the observed enlargement of vessels under perfusion (black traces Fig. 7B). SVideo 1 show changes in microtissue morphology under low FR (∼200µl/min, like that used in fibrin systems) suggesting that at this FR the tissue is perfused only during the peak of pressure due to the peristaltic cycle of the pump and that elastic return of the microtissue might cause oscillating flow in the capillaries. Higher FR (> 400µL/min) did not produce this effect, with the medium flowing linearly through the network without evident oscillations.

## Discussion

Creating biomimetic, tissue specific and vascularised microenvironments is a current challenge in tissue engineering. Overcoming this challenge can help addressing greater ones in basic cell biology, regenerative medicine and cancer research. Examples span from the potential to generate functional stem cells derived microtissues *in vitro* for regenerative medicine, or to create predictive disease models (e.g., cancer-on-chip) for drug testing and development. Overall, creating healthy and pathologic microtissues *in vitro* from primary cells or stem cells is revolutionising the way we conduct basic investigations in health and disease and creating appropriate vasculature in these tissues is a major bottleneck to further progression. Here we have addressed this challenge by creating a perfusable, scalable, customisable, and low-cost Vessel on Chip (VoC) platform.

Barriers to adoption of microfluidic-based lab on chip (LoC) are high as commercially available LoC and perfusion drivers are usually expensive and relatively fixed in design. In this work, we have demonstrated a workflow to fabricate inexpensive, custom and scalable LoC devices. The workflow in the current research is based upon 3DP and includes validation steps to engineer and manufacture low-cost LoC (Fig. 1) which can be rapidly modified and tailored to specific applications. One of the major challenges in the design of LoC/VoC systems is introducing continuous and controlled perfusion. By integrating a low-cost and compact flow driver in our design and by using 3DP to facilitate assembly, we report the creation and validation of a perfusion system that can be integrated in any lab with standard cell culture incubators (Fig. 2).

To accelerate LoC system design, we leveraged on the capabilities of Computational Flow Dynamics (CFD) as preliminary validation (Fig. 3) which has been invaluable during the development of our VoC prototype. CFD allowed us to conduct a preliminary *in silico* screening of possible designs and to rule out non-viable candidates. Furthermore, CFD allowed us to fine tune many parameters of our VoC design and optimise it for both functionality and easy assembly. Overall, the use of CFD enormously sped up and reduced the costs of VoC prototype development. As shown in the CFD simulations (Fig. 3), different compartments receive distinct flow, experiencing a net pressure drop between the aA and eV sides, essential for the creation of functional and hierarchical vasculature. Our design allowed creating laminar flow into the central well to reduce biomechanical stresses in the first days of perfusion when the vasculature is still fragile. Overall, we believe that parallel integration of 3D CAD, 3DP and CFD offers great potential and will be of particular interest for future implementation of multi-organ chips and the study of vascular dynamics in engineered vascular beds.

For initial validation of our VoC we used a fibrin gel vasculogenesis assay and shown that our VoC and perfusion system enables continuous long term (>15 days) perfusion of engineered vascular networks. However, vascular networks formed in hydrogels cannot recapitulate vascular maturation because the cellular components enabling matrix remodelling (e.g., macrophages and matrix producing stromal cells) are absent in such systems. To develop a more biomimetic model of microvasculature *in vitro*, we adapted an organotypic co-culture system of matrix secreting stromal and endothelial cells (OVAA, Fig. 5). OVAA exploits the ability of stromal cells like HDF to secrete the ECM necessary to embed ECs and form a capillary network with patent microvessels resembling *in vivo* counterparts. We have shown that OVAA can be used with different EC and stromal cells enabling the creation of tissue specific microenvironments (Fig. 5 and SFig. 2C).

Finally, we combined and optimised our platforms to create an integrated VoC-OVAA platform for the perfusion of microtissues. After 7-10 days of culture, ECs from the side channels and the central well assembled into a continuous vascular tree that could be perfused (Fig. 6). To the best of our knowledge, this is the first reported platform enabling continuous and long-term perfusion of vascularised tissues (Figs. 4, 6 and 7). A key advantage of VoC-OVAA is the physiological biomechanical forces and flow experienced by the cell as observed in Fig. 7 and SVideo. 1. As reported in Fig. 7, continuously perfused capillary networks remodel in response to these forces over time. Therefore, our platform paves the way to generating tissue specific microenvironments surpassing current hydrogel-based strategies and allows creating vascularised tissues which are easy to image, possess physiological biomechanical properties and are highly customisable.

Our VoC-OVAA will enable investigating a variety of key mechanisms in vascular biology which have physio pathologic implications. For example, cancer-associated signalling and extracellular matrix stiffening has been associated with impaired vascular maturation and reduced oxygenation of cancer tissue (Wu et al., 2021). Vascular abnormalities are also associated with impaired delivery of chemotherapies and immunotherapies with overall worse prognosis, and vascular normalisation therapy represents a very promising avenue to improve therapy outcome (Jain, 2005). Our VoC-OVAA system is by design able to model several of these aspects allowing to investigate in fine detail the molecular dynamics underpinning physiologic or impaired vascular maturation opening the way to more precise drug targeting. In this sense, another interesting potential application of our VoC-OVAA is extensive culture of tissue slice which can be derived from healthy and pathologic human tissue (Nogueira et al., 2022; Pitoulis et al., 2020) cultured into a VoC and used for example for drug sensitivity assays. Our VoC-OVAA can be easily adapted to create interconnected multi-organ chips by creating multichambered VoC each containing separate microenvironments interconnected by a common circulation system.

Finally, the VoC-OVAA platform presented here opens the way to develop models of vascularised organoids of stem cell-based systems where stem cell derived EC, perivascular and stromal cells as well as tissue parenchyma can be cultured or co/differentiated to create functional microtissues. Such pursuit will not only immensely improve our knowledge over cellular and molecular mechanisms of vasculogenesis, angiogenesis and organogenesis but also deliver new animal-free platforms for regenerative medicine, disease modelling and drug discovery.

## Materials and methods

### Computer assisted design (CAD)

The LoC device moulds, the plate containing the LoC device and the cassette for the microfluidic pump were designed using AutoDesk Fusion 360 (Autodesk, California, USA). Once created, the 3D objects were exported in STL format and sent to print on the different 3D printers or used for Computational Flow Dynamics (CFD) experiments.

### Computational flow dynamics simulations

All CFD simulations were performed using SimScale (SimScale GmbH, Germany). For all simulations we employed the incompressible flow formalism including the k-omega SST turbulence model, which, allows estimating the dynamics of medium flow through our chips. The parameters used where seawater 3.5 pc saline for the material of choice, 100 or 1000 µl/min for the velocity inlet and outlet values and mm for the model units. At the scales (channels sections 100-250×100-250 µm) and flow rates (FR) considered flow of medium containing 10% serum is in principle laminar throughout the chip (Reynolds number <10^2^ estimated by considering linear square section pipes with side length = 100-250 µm and 100 or 1000 µl/min fixed flow rate). CFD results were visualised using the Simscale data post-processing tool including cut-plane, particle trace analyses and corresponding statistics. We repeated all simulations at least three times and all our simulation converged to stable flow after 500s (all simulations were run for 1000s with 1s steps).

### Mould printing via stereolithography and post-processing

The mould were printed using UV-curable resin with a DLP (Digital Light Processing) 3D printer (MiiCraft ultra series 125), using transparent resin BV-007. After printing, the mould is detached from the platform and washed extensively to remove remaining resin. Briefly, it is rinsed in isopropanol, then sonicated twice in isopropanol for 5 minutes. The mould is then sonicated 5 minutes in acetone before being dried and post-cured in a UV chamber for 1h at 60°C. This process is repeated twice, followed by vapour phase deposition of Trichloro (1H, 1H, 2H, 2H-perfluroctyl) silane 97% in a desiccator for 60 minutes.

### PDMS preparation, tube embedding and casting

The LoC devices are manufactured by soft lithography on 3D printed moulds using silicon elastomer (Sylgard 184, Corning) and fixed on glass slides including silicon tubing (1.5mm ID, sourcing map) for world-to-LoC-connections.For the creation of the LoC device, polydimethylsiloxane (PDMS, SylgardTM 184 Silicone Elastomer) cast is prepared at a 1:10 base/curing agent ratio. The mixture is poured on the mould after placing the silicone tubing on the pillars to embed the tubing in the LoC device. Bubbles are removed from the silicone by a vacuum system before the mould is put to cure in the oven at 60°C for minimum 3h.

### LoC assembly and sterilisation

Once the silicone is cured, the PDMS can be peeled off the mould using the walls and cut using cutting guides to create individual LoC devices. LoCs are bonded to a glass slide, upon activation of the PDMS and glass surface via plasma treatment with a corona discharge device (piezobrush^®^ PZ2, Relyon plasma, Germany). To create a medium reservoir, rectangular walls of ABS (Acrylonitrile butadiene styrene) are printed with a FDM 3D printer (Cubicon Single Plus, Cubicon, Republic of Korea) and glued on top of the PDMS with PDMS for 1h in the oven. After ensuring that the LoC is properly assembled and sealed, the chip is autoclaved for sterilisation.

### 3D printed cassette with fused deposition modelling

The LoC device cassette is printed in ABS (Cubicon Single Plus, Cubicon, Republic of Korea). It is designed to keep the LoC device sterile therefore has standard cell culture plate dimension and can fit a cell culture plate lid for imaging, as well as transparent bottom, wells to create a humidity chamber and slots to host the microfluidic tubing.

### Microfluidic pump assembly and system sterilisation

After the LoC is assembled and sterile, it can be used to culture cells in a normal cell culture petri dish or the 3DP cassette and linked to the microfluidic system. For the culture of ECs, the chip is coated with bovine fibronectin solution (1mg/mL, Promocell) for at least 3h before cell seeding.

The microfluidic system used was the mpSmart-Lowdosing (Bartels mikrotechnik, Germany) including a microfluidic peristaltic pump (0.005-1mL/min), a filter (average 0.45µm pore size) and a flow sensor (Sensirion SLF3s-0600F) linked to a laptop via a controller driver for controllable FR. Before LoC perfusion, the system including tubing is sterilised by perfusing 1% sodium hypochlorite for 10min then rinsed thoroughly with distilled water.

### Cell culture

Human Umbilical Vein Endothelial Cells (HUVEC, PromoCell), Human Aortic Endothelial Cells (HAoEC, Promocell), Human Pulmonary Microvascular Endothelial Cells (HPMEC, Promocell) and Red Fluorescent Protein-tagged HUVEC (RFP-HUVEC, 2Bscientific, UK) were cultured on fibronectin-coated (PromoCell) plates up to passage 7 and maintained in Endothelial Growth Medium 2 (EGM2; Promocell). Normal Human Dermal Fibroblasts (NHDF, Promocell) were cultured in Fibroblast Growth Medium 2 (Promocell) up to passage 7. Human Mesenchymal Stem Cells (MSC, PromoCell) were cultured in Mesenchymal Growth medium 2 (PromoCell) up to passage 7. Cells were routinely passaged using Accutase™ (Thermo). The cell line THP-1 (ATCC, UK) was cultured in suspension in DMEM (Gibco) supplemented with 10% FBS (Thermo).

### Fibrin gel vasculogenesis assay

The fibrin gel was formed by mixing an 8 mg/mL solution of fibrinogen (Sigma) containing 15.10^6^ EC/mL and a 4 U/mL thrombin solution (Sigma) and 10μM Aprotinin (Thermo) in EGM2 medium (Promocell). 20 μL of the mixture was pipetted in the central well of the LoC device and left 30 min at 37 °C for hydrogel setting, before the reservoir and channels were filled with medium. The chip was cultured in a petri dish in presence of distilled water to create a humidity chamber and prevent medium evaporation before connection to the microfluidic system in the 3D-printed cassette.

### OVAA in static conditions

The method used is an optimised version of previously published one (Hetheridge et al., 2011). Briefly, 7.5×10^3^/cm^2^ Red Fluorescent Protein-tagged HUVEC (RFP-HUVEC, 2Bscientific, UK) are mixed and seeded with 1.5×10^5^/cm^2^ Normal Human Dermal Fibroblasts (NHDF, Promocell) and left in culture for 14 days to allow EC vasculogenesis with medium change (supplemented or not with 50ng/mL VEGFA, Peprotech, UK) every two days. Cultures are monitored daily to assess generation of microvascular structures which start to form around day 5 and can be maintained in culture for more than 30 days under static conditions. After 14 days of culture, cells are fixed and immunostained for CD31 (ab9498, 1μg/mL, Abcam) and revealed with fluorescently labelled secondary antibody goat anti-mouse (A-21127, 1μg/mL, ThermoFisher). Images were obtained using an Operetta CLS system (PerkinElmer, Waltham, MA) equipped with a 10X objective for full well scans and 40X and 63X immersion objective for high resolution images. Images were quantified using the ImageJ AngioAnalyser plugin (Carpentier et al., 2020).

### Vasculature-on-chip culture and continuous perfusion

The VoC is first cultured in static conditions in a 75cm^2^ petri dish (Greiner bio-one) before perfusion in the 3D-printed cassette. HUVEC-RFP are first cultured in monolayers in the LoC device in the channels after fibronectin coating (PromoCell) for 3 to 5 days. Then, a co-culture of 7.5×10^3^ Red Fluorescent Protein-tagged HUVEC (RFP-HUVEC, 2Bscientific, UK) and 1×10^5^ Normal Human Dermal Fibroblasts (NHDF, Promocell) or Human Mesenchymal Stem cells (MSC, PromoCell) are seeded in the central well of the device. Cells are kept in static culture for 7 days and maintained in EGMV2 (PromoCell) changed every other day from the top of the device. After monitoring of vascular network formation, the device is transferred in the 3D-printed cassette and linked to the microfluidic pump device to be perfused continuously. Images from the vascular network and videos of the perfusion were obtained with a wide field inverted microscope (Olympus IX51, Biosystems, Munich, Germany). Confocal images were obtained with a Leica Sp8 confocal inverted microscope, from the RFP signal of the HUVEC-RFP or, after fixation, from the EC-specific VE-cadherin signal (Novusbio NB600-1409, 1μg/mL) counter-stained with a secondary Alexa Fluor 488 (A-21141, 1μg/mL, Thermo Fisher Scientific).

## Acknowledgments

The authors wish to thank the following people/organisations which offered their help and assistance at various phases during the development of this project. iMakr (London, UK) for lending the Miicraft 3D printer and offering their expertise in 3D printing during early phases of the project. Kerushan Thomas, iBSC TEIT student 2019-2020 for helping to develop the design of the LWC. Florian Siemenroth and Bartels Mikrotechnik GmbH (Dortmund, GE) for their assistance in developing a custom microfluidic driver system.

## Authors Contributions

Conceptualization: FC, LV; Methodology: FC, JJ, JH, SS, LV; Formal analysis: FC, LV; Writing: FC, LV; Supervision and Advice: LDS, AG, TC, LV; Critical review of the manuscript: LDS, AG, TC; Project administration: LV

## Captions to figures

<u>Supplementary figure 1: LoC device design optimisation.</u>

A) Different design of the side channel length and configuration. Designs that were considered include: independent (i-ii) afferent arteriole (aA) and efferent venule (eV), direct total perfusion (iii) in the central well, or partially divided (iv, v) flow into the central well. B) The junction between aA and eV can be designed to have a 90° angle (i), a bigger angle (ii) to receive more flow or a longer length (iii).

<u>Supplementary figure 2: VoC timeline and versatility.</u>

A) Timeline for the creation of perfused vasculature-on-chip. The VoC is first coated with fibronectin to promote the adhesion of ECs in the side channels. At day 0, ECs are seeded in the side channels (aA and eV) and left in culture for 5 days to coat the channels on all sides. At day 5, the co-culture of ECs and FBs is seeded in the central well and cultured in static conditions for 7 days to promote capillary network formation. At day 12, the VoC can be linked to the microfluidic system and continuously perfused. B) Photographs of the VoC system with HUVEC-RFP at day 0, 5 and 12 of culture in co-culture with HDF. Scale bar: 500μm. C) Photographs of the VoC system with HUVEC-RFP at day 0 and 12 of culture in co-culture with MSCs. Scale bar: 250μm.

<u>Supplementary figure 3</u>: 3D projection of the vascular network formed by HUVEC-RFP in the central well of the VoC after 7 days of culture, before perfusion.

<u>Supplementary video 1</u>: Video of the VoC system at one of the aA junctions after 4 days of perfusion at 200uL/min.

