## Supplementary figures and images for "Continuously perfusable, customisable and matrix-free vasculature on a chip platform"

### Supplementary figure 1

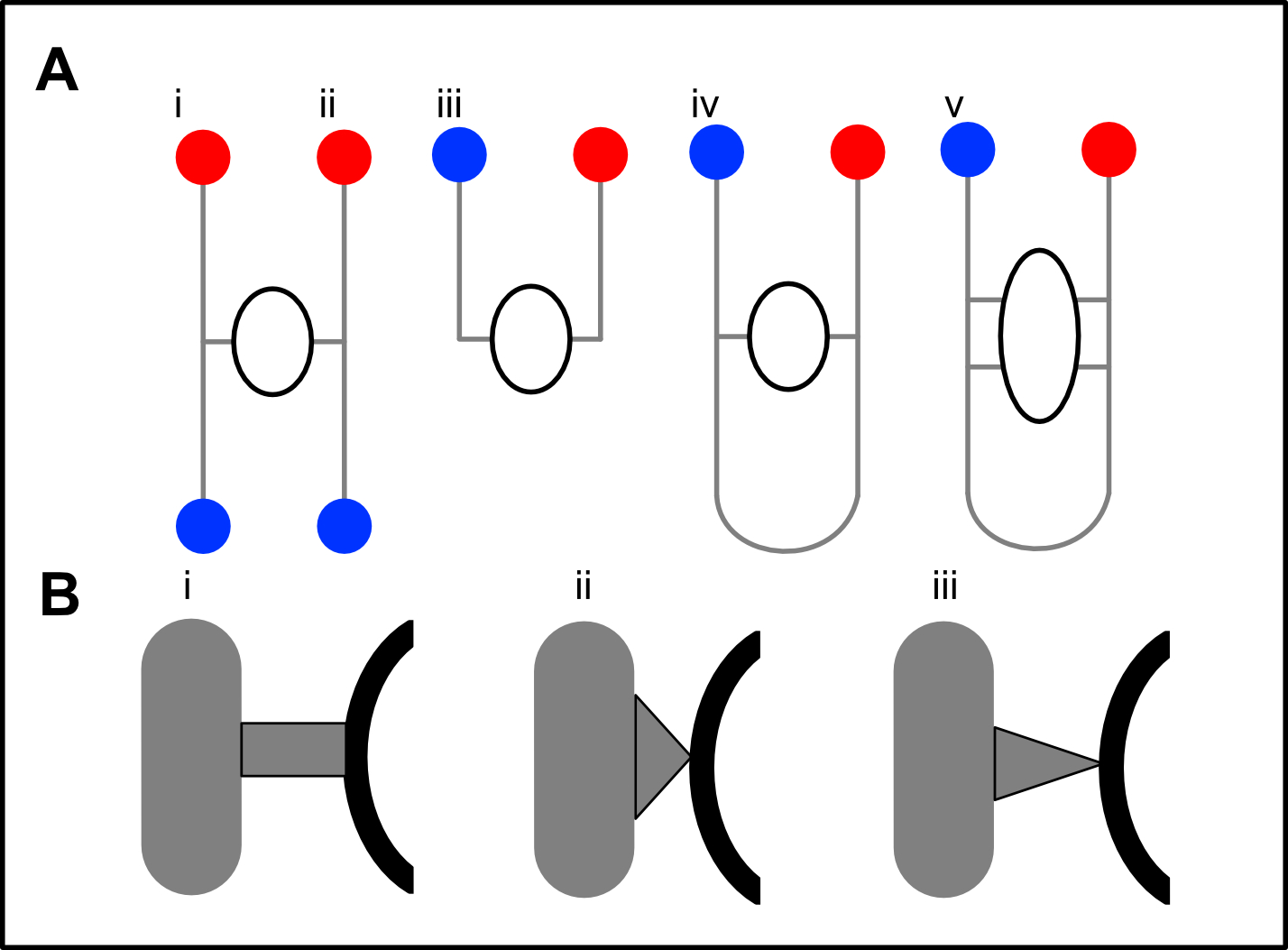

### Supplementary figure 2

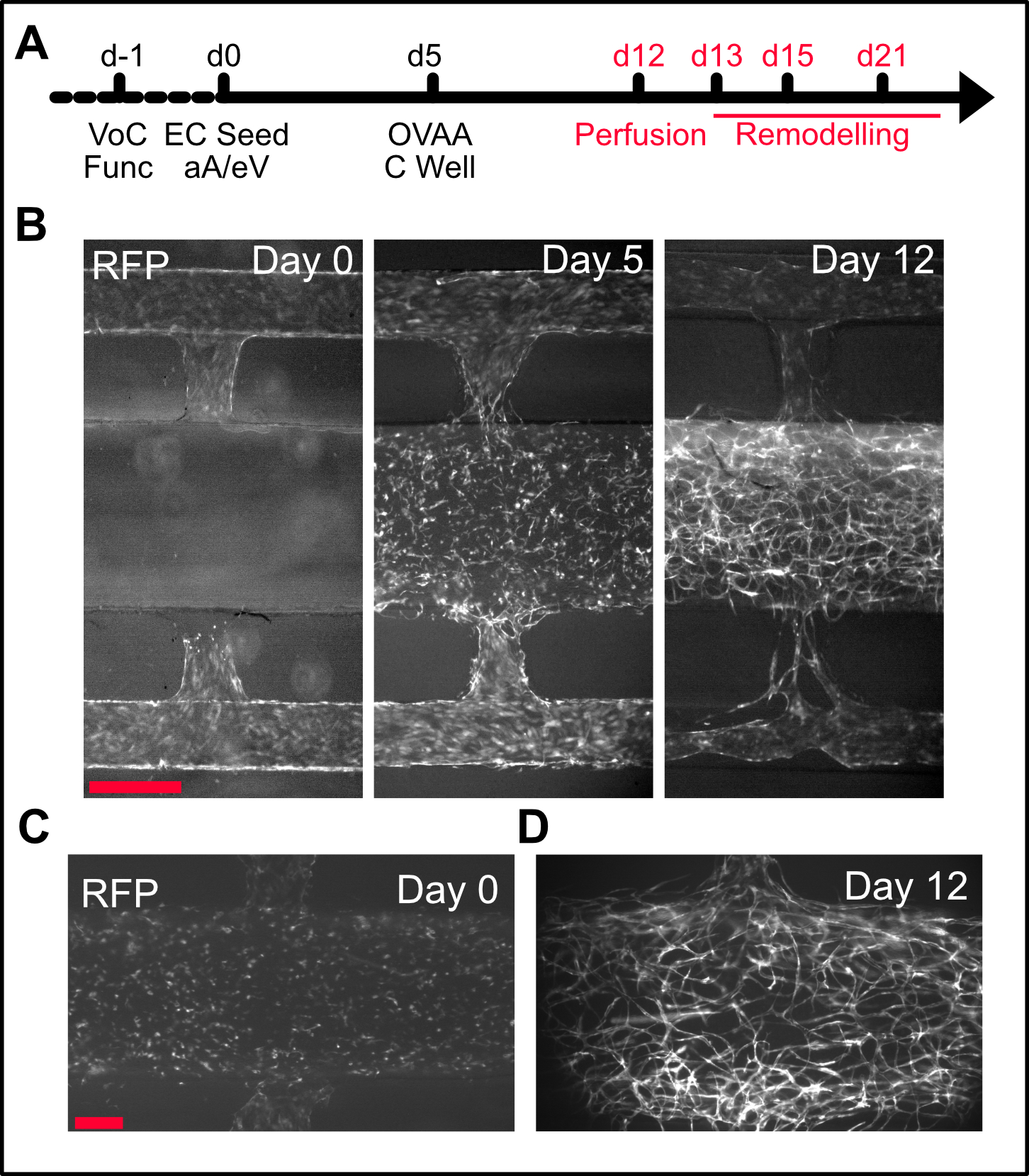
